# Real-time fMRI neurofeedback reduces default mode network and auditory cortex functional connectivity in schizophrenia

**DOI:** 10.1101/2025.01.02.631107

**Authors:** Jiahe Zhang, Clemens C.C. Bauer, Francesca Morfini, Yoonji Lee, Lena Stone, Angelina Awad, Kana Okano, Melissa Hwang, Ann K. Shinn, Margaret A. Niznikiewicz, Susan Whitfield-Gabrieli

## Abstract

**Background and Hypothesis:** Auditory verbal hallucinations (AHs) are a cardinal symptom of schizophrenia that can cause distress but are not always responsive to antipsychotic medications. There is a critical need to develop novel interventions that target neural mechanisms underlying AHs. We developed a real-time fMRI neurofeedback (NFB) paradigm for AHs that aims at modulating default mode network (DMN) functional connectivity.

**Study Design:** Patients with schizophrenia or schizoaffective disorders who were experiencing AHs (*N* = 25) attempted to decrease brain activation while listening to sentences recorded in another person’s voice and increase brain activation while listening to sentences recorded in their own voice. Participants randomly assigned to the ‘real’ group (*n* = 12) received neurofeedback based on signals from their auditory cortex in the superior temporal gyrus (STG) and those assigned to the ‘sham’ group (*n* = 13) received neurofeedback based on motor cortex signals.

**Study Results:** Analyzing resting state fMRI data collected pre- and post-NFB, we found that: (1) at baseline, stronger within-DMN connectivity between the medial prefrontal cortex (MPFC) and posterior cingulate cortex was associated with higher AHs severity; (2) after NFB, participants in the real group, compared to those in the sham group, showed greater reduction in functional connectivity between the MPFC and auditory cortices in the STG and middle temporal gyrus (MTG). Notably, the reduction in MPFC-STG/MTG connectivity was found in all participants in the real group.

**Conclusions:** These findings suggest that NFB can effectively and non-invasively modulate functional connectivity in regions associated with AHs in psychosis.

## INTRODUCTION

Auditory verbal hallucinations (AHs) affect over 70% of patients with schizophrenia (SZ)^1–6^ and one third of those suffering from AHs do not respond to antipsychotic medications,^3,7^ underscoring the urgent need to develop interventions that target the core mechanisms of AHs in SZ.

Recent neuroimaging studies reveal that SZ is associated with disruptions in large-scale neural circuits, such as the default mode network (DMN), which typically activates during self-referential processing and spontaneous thought.^8,9^ Compared to healthy controls, patients with, or at risk for, SZ exhibit increased DMN functional connectivity between its midline hubs of medial prefrontal cortex (MPFC) and posterior cingulate cortex (PCC),^10–13^ suggesting impaired self- and reality-monitoring.^14^ Studies on AHs further implicate the auditory cortices (i.e., superior and middle temporal gyri) based on their heightened activity before the onset of and during auditory hallucinations^15^ and their altered functional connectivity with the DMN.^16–18^ It has been hypothesized that abnormally increased connectivity between the MPFC and auditory cortices in the superior temporal gyrus (STG) may contribute to source misattribution of self-generated auditory content as external stimuli.^19^

Given the association between increased STG activity and AHs, we developed a novel, non-invasive intervention that combines mindfulness with real-time functional magnetic resonance imaging (fMRI) neurofeedback (NFB) to help individuals to volitionally downregulate STG activity.^20,21^ In this intervention, individuals observe a visual representation of their STG activity to evaluate whether their effort to downregulate their STG activity, aided by mindfulness practice, leads to the expected reduction. Individuals use this neurofeedback to modify their effort during the next trial. We have previously shown that participants receiving STG-based NFB successfully reduced STG activation in SZ patients with AHs.^20^ Other NFB studies based on network-level activation have also shown evidence of network-level functional connectivity changes such as reduced DMN connectivity in SZ with AHs^21^ and DMN-STG connectivity changes in SZ with AHs.^22^ However, these NFB studies did not include a sham control condition. Furthermore, it remains unclear if STG-based NFB, which is based on neurofeedback from a single region, can similarly lead to functional connectivity changes in associated brain networks such as the DMN.

In the current randomized controlled trial, patients with a SZ spectrum disorder and moderate to severe AHs were randomized to receive NFB from either STG activity (‘real’ group) or motor cortex activity (‘sham’ group). Other analyses based on this research project (clinicaltrials.gov: NCT03504579) have reported reduced STG activation while listening to sentences recorded in the participant’s own voice^23^ and increased DMN activation during self-referential processing.^24^ For the current paper, we collected resting state fMRI scans before and after NFB to examine 1) baseline correlations between DMN/STG functional connectivity and AHs severity, and 2) the effect of NFB on DMN/STG functional connectivity.

## METHODS

### Participants and Procedures

Forty-one patients were enrolled in this study. Eligibility criteria included i) a diagnosis of schizophrenia or schizoaffective disorder verified with the Structured Clinical Interview for DSM-5 (SCID-5) interview; ii) moderate or severe medication-resistant AHs (≥ 4 on item #3 of the Positive and Negative Syndrome Scale (PANSS) in the past month)); iii) 18-55 years old; iv) on stable doses of medications; v) right-handedness (Edinburgh Handedness Inventory ≥ 60); vi) native English speakers; vii) estimated IQ ≥ 70 (WASI); and viii) normal or corrected-to-normal vision and normal hearing.

Participants were excluded if they i) had a history of neurological or traumatic head injury in the past six months; ii) had alcohol use disorder or substance use disorder in the past month (as per DSM-5) or used alcohol in the previous 24 hours; iii) were pregnant; or iv) had other MRI contraindications.

All participants gave written consent obtained in accordance with the guidelines by Institutional Review Boards of Harvard Medical School, Boston VA Healthcare System, Massachusetts Institute of Technology (MIT), and Northeastern University. As we have previously described,^24^ participants underwent four sessions including an initial clinical visit, a baseline fMRI visit for functional localization, a NFB fMRI session, and a post-NFB follow-up visit. Details about each session are below.

Six participants were deemed ineligible due to exclusion criteria after enrollment. Five participants withdrew during the course of the study. Four participants did not have complete data as they were lost to contact during the course of the study. This resulted in a final sample of 25 participants (36.1 ± 10.0 years; 24-54 years; 24% females) who were randomly assigned to receive either real neurofeedback (*n* = 12) or sham neurofeedback (*n* = 13). Two participants were lost to contact following NFB session and therefore had missing post-NFB AHs scores. Another two participants (one in each condition) completed follow-up session more than 200 days after NFB session due to COVID lockdown, therefore we analyzed their rsfMRI data from baseline and NFB fMRI visits but excluded their post-NFB AHs scores from further analyses. See more details in the CONSORT diagram in **Fig. S1**.

During the course of our study, we were required to change sites and scanners. Among the 25 participants, seven were scanned on a Siemens Trio scanner at MIT, four were scanned on a Siemens Prisma scanner at MIT, and 14 were scanned on a Siemens Prisma scanner at Northeastern University.

We administered the auditory hallucinations subscale of the Psychotic Symptom Rating Scales (PSYRATS-AH)^25^ to measure AHs severity. The PSYRATS-AH measures 11 items about AHs: frequency, duration, location, loudness, beliefs regarding the origin of voices, amount of negative content or voices, degree of negative content, amount and intensity of distress, disruption to life caused by voices, and controllability Each item is rated on a scale of 0 -to 4 for a total PSYRATS-AH score ranging from 0-44 points, where higher PSYRATS scores indicate more severe AHs.

### Clinical visit

After obtaining informed consent, we collected participants’ demographic and clinical information. We administered the SCID-5 to determine psychiatric diagnoses, the PANSS to assess psychosis severity, and the PSYRATS-AH to measure AHs severity. We also conducted neuropsychological testing (data not presented here). Finally, we obtained audio recordings while participants read a series of sentences out loud, to be used during subsequent fMRI sessions.

### Baseline fMRI visit

Participants were administered the PSYRATS-AH at the beginning of the session, then underwent functional localization procedures in the scanner (see **Supplementary Materials** for MRI acquisition details). In the current study, two functional localizers were performed to facilitate identification of participant-specific brain regions whose blood-oxygen level dependent (BOLD) fluctuations serve as the source of neurofeedback signals.

#### STG Functional Localizer

For STG functional localization, participants completed a self-other-voice task in which they passively listened to a series of sentences pre-recorded in their own voice (self condition), in a stranger’s voice (other condition), or while fixated on a crosshair (silence condition). Two 4.5-minute runs were acquired, each including a 30-second baseline and twelve 16-second blocks (four blocks per condition). Each block for self or other condition included 5 unique sentences. Preprocessing of fMRI data was performed in FSL 6.0^26^ and included: realignment, skull stripping, co-registration, smoothing with 5mm Gaussian kernel and high-pass filter (100 seconds). We defined individualized STG as the clusters with the maximum voxel intensity for self>other blocks within bilateral anterior and posterior STG regions of the Harvard-Oxford 25% probability atlas.

#### Motor Cortex (MC) Functional Localizer

We localized each participant’s MC using resting state fMRI (rs-fMRI). Preprocessing of rs-fMRI data was performed in FSL 6.0 ^26^ and included: motion correction, brain extraction, co-registration, smoothing and bandpass filtering (see more details in ^21^). We performed an independent components analysis (ICA) on the preprocessed functional scans using Melodic ICA version 3.14^27^ with dimensionality estimation using the Laplace approximation to the Bayesian evidence of the model. We extracted 30 spatiotemporal components and compared them to an atlas map of the motor network.^28^ We used FSL’s *fslcc* tool and selected the ICA components that yielded the highest spatial correlation for each participant. These ICA components were thresholded to select the upper 10% of voxel loadings and then binarized to obtain participant-specific MC masks. Visual inspection of the components was performed to ensure satisfactory coverage of the canonical MC.^29^

### Real-time fMRI neurofeedback (NFB) visit

NFB session occurred an average of 10.0 ± 10.2 days after baseline fMRI session. In NFB session, participants completed: mindfulness training, pre-NFB rs-fMRI, NFB, post-NFB rs-fMRI.

#### Mindfulness Training

We trained participants on a mindfulness technique called “mental noting”. The experimenter explained to each participant that mental noting entails being aware of a sensory experience without engaging in or dwelling on the details of the content; in other words, one would “note” the sensory modality (e.g., “hearing”, “seeing”, “feeling”) at the forefront of their awareness and then let go of the experience after it has been noted. The experimenter also introduced the concept of an “anchor”, or a sensory experience to which one could easily switch their attention, such as breathing.

Participants were encouraged to use their personal “anchors” when they made consecutive notes of “hearing”. The experimenter demonstrated noting practice by verbalizing out loud the predominant sensory modality approximately once per second. The participant then practiced mental noting out loud. Prior to proceeding to the NFB task, the experimenter ensured the participant described sensory awareness without engaging in the content and stopped consecutive “hearing”.

#### NFB Task

During the neurofeedback task, an analysis computer continuously received fMRI data from the MRI scanner and computed the averaged activation level within either the individualized STG or MC masks based on the raw data in real time. Averaged activation during a task block was visually displayed to the participant as a ‘thermometer’ on a projection screen. (**Fig. 1A**). The neurofeedback task followed a blocked design with two conditions (**Fig. 1B**). In the ‘Listen’ condition, participants listened to five sentences in their own voice and were instructed to attend to the auditory stimulus. In the ‘Ignore’ condition, participants listened to five sentences in a sex-matched other voice and were instructed to use mindfulness to ignore the auditory stimulus. After each block of five sentences, participants were asked to assess their own performance via button press and then were presented with the image of a thermometer displaying neurofeedback based on real-time fMRI analysis of their ability to modulate STG activity during that block. There were four neurofeedback runs in this protocol and each lasted 2.5 minutes.

**Figure 1.**
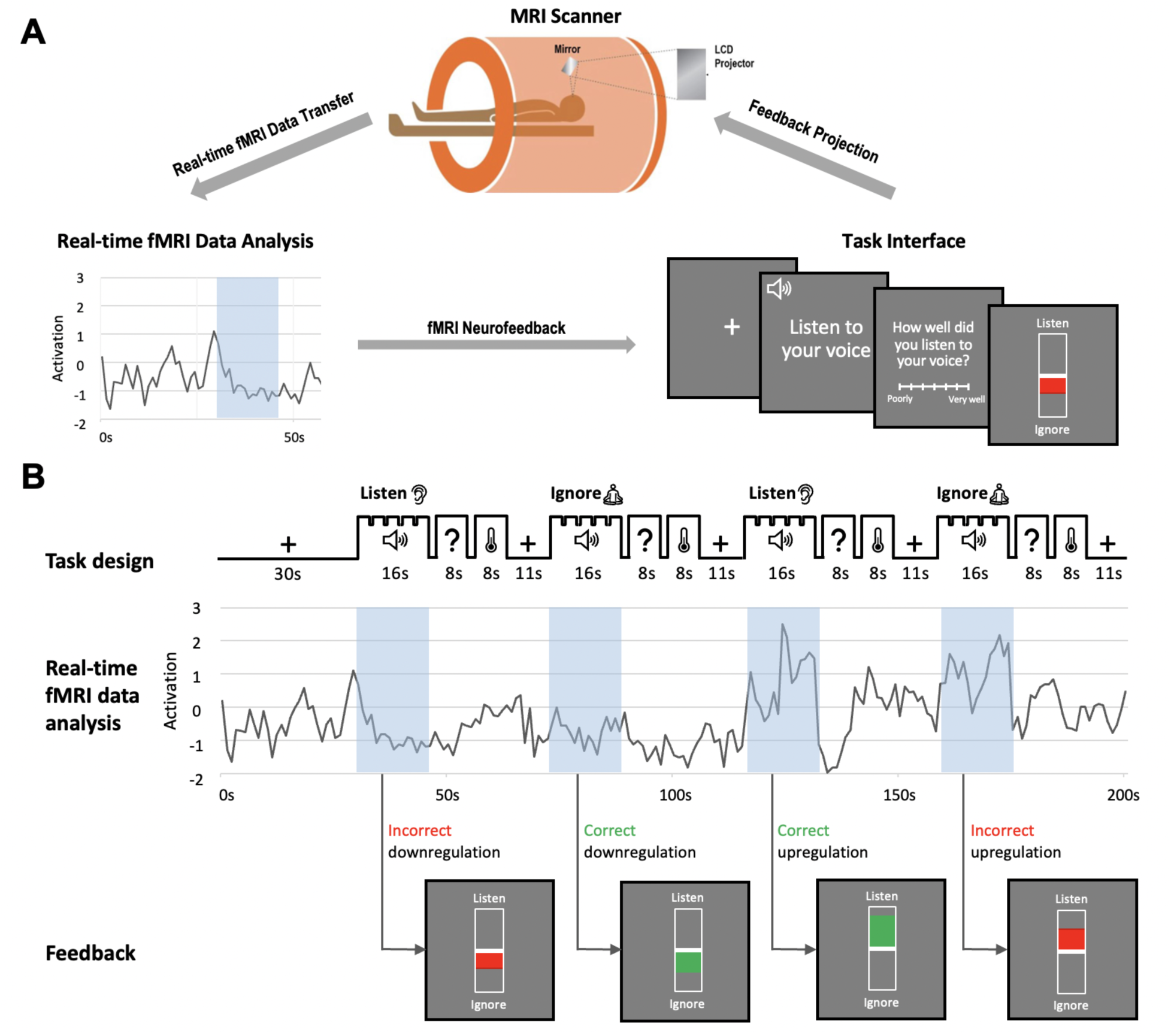
Mindfulness-based real-time fMRI neurofeedback experiment. **A)** The real-time fMRI neurofeedback setup includes three main components that form a loop of information flow. **B)** A simulated fMRI time series illustrates four possible types of neurofeedback that a participant could receive. Green color on the thermometer indicates modulation in the correct direction (i.e., downregulation during ‘Ignore’ condition or upregulation during ‘Listen’ condition) and red color indicates modulation in the wrong direction (i.e., downregulation during ‘Listen’ condition or upregulation during ‘Ignore’ condition).

#### NFB Calculation

We provided real-time neurofeedback of STG activation for the real group and MC activation for the sham group. Activation estimates were calculated as the mean activity across all voxels within each participant’s STG mask (as defined by *STG Functional Localizer*) or MC mask (as defined by *MC Functional Localizer*) and co-registered to the current fMRI volumes. We used the signal from the MC in the sham condition since signal fluctuations in the MC are likely unrelated to hallucinations.^14,30^ To accomplish the voxel-wise estimation in real-time,^31^ we first collected 30 seconds of baseline data and then continuously performed an incremental general linear model (GLM) fit with subsequent incoming images. This method accounts for the mean voxel signal and linear trends. To discount components of the voxel signal due to nuisance sources (e.g., low-frequency signal drifts), the GLM reconstruction of the expected voxel intensity at time *t* was subtracted from the measured voxel intensity at time *t*, leaving a residual signal that has components due to two sources: BOLD signal fluctuations and unmodeled fMRI noise. This residual was scaled by an estimate of voxel reliability, which was computed as the average GLM residual over the first 25 functional images of the baseline.

This analysis resulted in an estimate of the strength of activation at each voxel at time *t* in units of standard deviation.

### Follow-up visit

Post-NFB follow-up visit occurred an average of 33.9 ± 59.3 days after NFB session. Two participants (1 in each condition) had a long lapse between sessions (>200 days) due to lockdowns during the COVID-19 pandemic. Excluding these two participants, follow-up visits occurred 16.2 ± 10.2 days after NFB session.

#### AHs assessment

Participants were administered the PSYRATS-AH prior to undergoing a scan that is unrelated to the current analyses.

### Image analysis

#### MRI Preprocessing

Preprocessing was performed using *fMRIPrep* 21.0.0,^32,33^ which is based on *Nipype* 1.6.1.^34,35^ In short, common preprocessing steps were performed including realignment, co-registration, normalization, susceptibility distortion correction, segmentation of gray matter (GM), white matter (WM), cerebrospinal fluid (CSF) tissues, skull stripping, and confounds extraction. See **Supplementary Materials** for a detailed description. Visual quality control was performed on each preprocessed run.

Preprocessed data and confound time series were imported into the CONN Toolbox v20.b^36^ where outlier identification was performed with the Artifact Detection Tools (ART, www.nitrc.org/projects/artifact_detect). Volumes with global signal z > 5 and framewise displacement > 0.9mm compared to the previous frame were flagged as outliers. In addition, in-scanner mean motion was defined as the mean framewise displacement of the whole run^26^ and calculated separately for pre- and post-NFB rs-fMRI runs. Rs-fMRI runs were spatially smoothed with a 6mm Gaussian kernel. A principal component analysis identified noise components from the WM and CSF following CONN’s implementation^37^ of the aCompCorr method.^38^ During denoising, we regressed out the effect of the top 5 WM noise components, top 5 CSF noise components, 12 realignment parameters (3 translation, 3 rotation, and their first derivatives), linear drift and its first derivative effect, motion outliers, and applied a band-pass filter of 0.008 – 0.09 HZ.

#### Functional Connectivity Analysis

Using the CONN toolbox,^36^ we performed functional connectivity analysis using the MPFC (MPFC node of the DMN network derived from an independent component analysis of the Human Connectome Project dataset, *N* = 497^37^) and the STG (8mm sphere around MNI 60, −18, 10^20^) as seed regions. For baseline brain-behavior correlation analysis across the full sample, we searched the whole brain for voxels whose connectivity with the seed correlated with PSYRATS-AH scores. For functional connectivity change, we searched the whole brain for voxels whose connectivity with the seed region changed significantly after NFB in the real group compared to the sham group. To specifically test for MPFC-PCC connectivity change in the real group, we used SPM small volume correction to search for PCC voxels whose connectivity with the MPFC seed region changed significantly after NFB and reported a cluster that survived a FDR-corrected threshold of *q*=.05. All analyses controlled for framewise displacement.^39^

## RESULTS

### Participants

Sociodemographic and clinical characteristics are summarized in **Table 1**. The real and sham groups did not differ on sociodemographic variables or in PSYRATS-AH scores pre- or post-NFB.

**Table 1.** Sociodemographic and clinical information.

|  | **Real** | **Sham** | **Real vs. Sham** |
| --- | --- | --- | --- |
| *N* | 12 | 13 |  |
| **Sociodemographic** |  |  |  |
| Age (mean/SD) | 35.9 (11.1) | 36.3 (9.1) | t-test: $p = 0.921$ |
| Sex: female ($n/\%$) | 16.7% | 30.8% | Wald test: $p = 0.409$ |
| Race: White ($n/\%$) | 58.3% | 84.6% | Wald test: $p = 0.144$ |
| Education (mean/SD) | 13.3 (2.4) | 13.8 (2.1) | t-test: $p = 0.545$ |
| IQ: WASI (mean/SD) | 100.7 (16.8) | 98.8 (13.3) | t-test: $p = 0.756$ |
| **PANSS (mean/SD)** |  |  |  |
| Total score | 67.7 (16.9) | 62.6 (14.8) | t-test: $p = 0.434$ |
| Positive subscale | 20.3 (7.5) | 19.0 (5.8) | t-test: $p = 0.599$ |
| Negative subscale | 15.3 (4.7) | 14.5 (4.7) | t-test: $p = 0.709$ |
| General subscale | 32.0 (8.7) | 29.1 (7.6) | t-test: $p = 0.378$ |
| **PSYRATS-AH (mean/SD)** |  |  |  |
| Baseline | 24.5 (7.5) | 25.2 (7.5) | t-test: $p = 0.809$ |
| Post-NFB | 18.5 (7.5) | 22.3 (7.3) | t-test: $p = 0.217$ |

*Note.* All real vs. sham tests were two-tailed independent samples tests. PANSS = Positive and Negative Symptoms Scale. PSYRATS-AH = Auditory Hallucination subscale of the Psychotic Symptom Rating Scales. WASI = Wechsler Abbreviated Scale Intelligence (Full Scale).

### Baseline DMN-AHs Symptom Severity Association

Baseline PSYRATS-AH positively correlated (p < 0.001, uncorrected) with functional connectivity between the MPFC seed (**Fig. 2A**) and a cluster in the PCC (**Fig. 2B**; 106 voxels; peak at MNI −10, −56, 22). Specifically, more severe AHs symptoms were associated with greater MPFC-PCC connectivity (**Fig. 2C**). Connectivity analysis with STG as the seed region yielded no significant correlations between baseline AHs severity and STG connectivity (*p* > 0.001, uncorrected).

**Figure 2.**
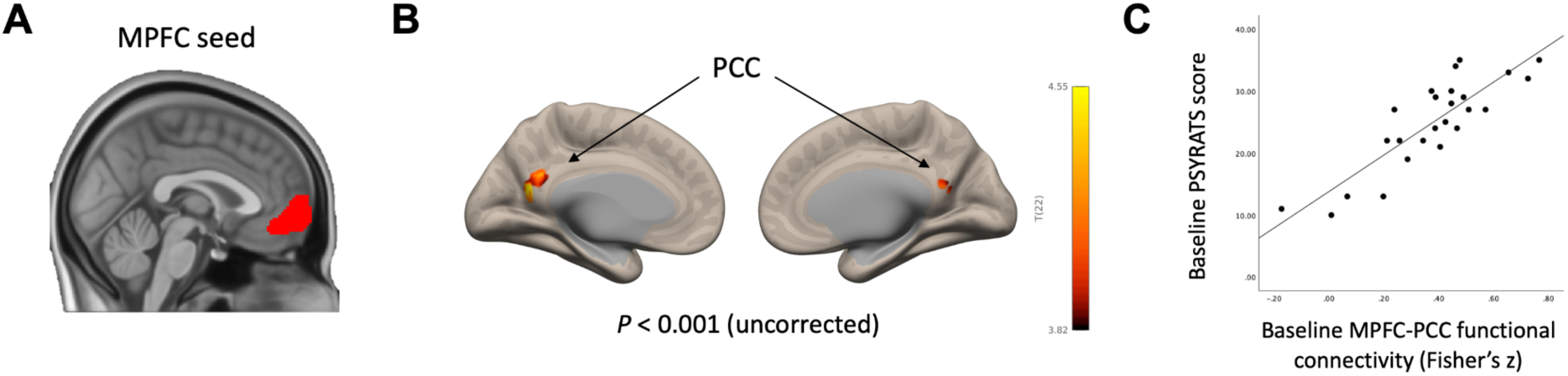
Higher DMN functional connectivity was associated with more severe auditory hallucination symptoms at baseline. **A)** We used the MPFC seed as part of the DMN network defined in the CONN toolbox ^37^. **B)** Functional connectivity between the MPFC seed and the PCC positively correlated with AHs severity (*p* < .001, uncorrected). Higher PSYRATS-AH score indicates higher severity. The color bar range reflects minimum and maximum *t* values in the connectivity map. **C)** Scatterplot illustrates the correlation between baseline PSYRATS-AH and baseline MPFC-PCC functional connectivity (computed using an averaged timecourse across all voxels in the significant PCC cluster).

### AHs Change Following NFB

A repeated measures ANOVA test revealed a significant main effect of time [F(1,19) = 6.66, *p* = 0.018], indicating significantly reduced AHs after NFB across both real and sham groups. The interaction between time and group was not significant [*F*(1,19) = 0.708, *p* = 0.411], indicating AHs change did not differ between real and sham groups (**Fig. S2**).

### Functional Connectivity Change Following NFB

Compared to the sham group, the real group showed significantly greater reductions in functional connectivity (*p* < 0.001, uncorrected) between the MPFC seed and clusters in bilateral auditory cortices (left MTG: 72 voxels, MNI −56, −28, −10; right STG/MTG: 147 voxels, MNI 60, −30, −4) and left inferior frontal gyrus (IFG: 76 voxels, MNI −56, 30, 2) (**Fig. 3A**; see connectivity maps for each group at each time point in **Fig. S3**). Notably, every participant in the real group (*n* = 12) showed a reduction in MPFC-STG/MTG connectivity, whereas almost all participants in the sham group (12 out of 13) showed MPFC-STG/MTG connectivity increase (**Fig. 3B**). Using an *a priori* STG seed that showed reduced activity post-NFB in a prior study (Okano et al,. 2020), we replicated the effect of significantly more reduced STG-MPFC connectivity in the real vs. sham group (**Fig. S4**).

**Figure 3.**
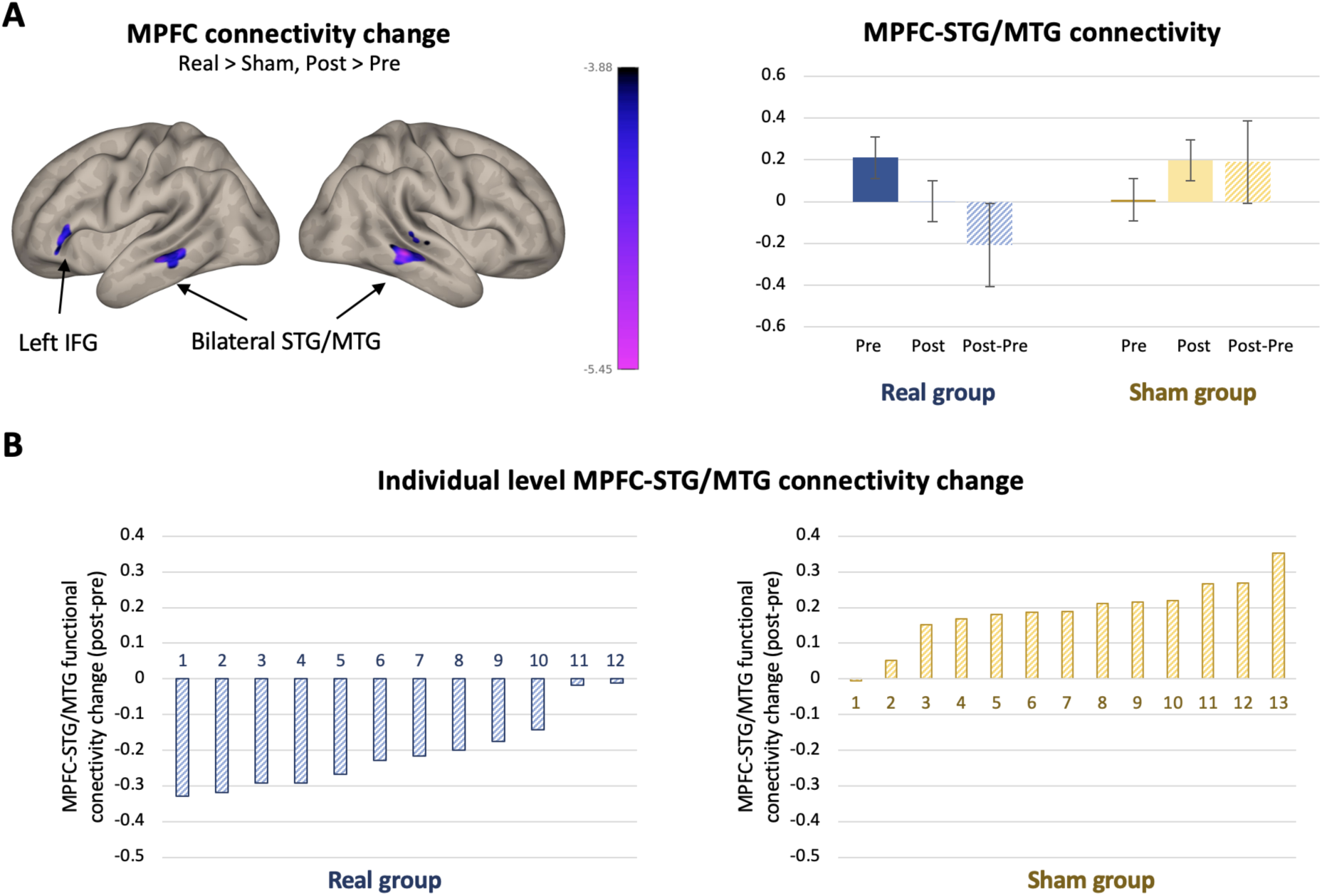
One session of NFB reduced MPFC-STG/MTG functional connectivity. **A)** An ANOVA test revealed that after NFB, there was significantly more reduced connectivity between MPFC seed and bilateral STG/MTG regions and left IFG (*p* < 0.001, uncorrected) in participants receiving real vs. sham NFB. Color bar ranges reflect minimum and maximum *t* values in the maps. The bar plot displays MPFC-STG/MTG connectivity (averaged between left and right STG/MTG clusters) pre-NFB and post-NFB, as well as their difference. Error bars indicate one standard error of the mean. **B)** Reduced MPFC-STG/MTG connectivity was found in all participants assigned to the real group (*n* = 12) and the opposite pattern of increased MPFC-STG/MTG connectivity was observed in all but one participant assigned to the sham group (*n* = 13). IFG: inferior frontal gyrus; MPFC: medial prefrontal cortex; MTG: middle temporal gyrus; STG: superior temporal gyrus.

Within the real group, a small volume correction analysis showed that MPFC connectivity with one PCC cluster (4833 voxels, MNI −4, −54, 36) changed significantly after NFB (*q*_FDR_ = 0.030) (**Fig. S5**). We extracted connectivity strength between the MPFC seed and the PCC cluster and ran a repeated measures ANOVA test, which revealed a marginally significant time-by-group interaction where participants in the real group showed more reduction in MPFC-PCC connectivity compared to the sham group [*F*(1,21) = 4.231, *p* = 0.052].

Within the real group or across the whole sample, AHs change did not correlate with MPFC-PCC or MPFC-STG/MTG connectivity change.

## DISCUSSION

AHs are an often debilitating symptom of SZ that are not always responsive to pharmacologic treatment. Real-time fMRI neurofeedback is a promising non-invasive alternative to traditional (e.g., medications, psychotherapy) and other emerging (e.g., transcranial magnetic stimulation) treatment approaches.^14,40,41^ Previous NFB studies targeting AHs explored the effects of downregulating the STG,^20,22,42^ anterior cingulate cortex,^43^ and the DMN.^21^ The current study adds several key findings to this emerging literature. First, with our baseline data, we replicated the finding of an association between stronger DMN connectivity and more severe SZ symptoms (i.e., AHs). Second, a major strength of our study was the introduction of a sham condition using neurofeedback from an alternative brain signal. We found that, compared to participants receiving sham MC-based neurofeedback, those receiving real STG-based neurofeedback showed more MPFC-STG/MTG connectivity reduction after NFB.

Our finding that stronger MPFC-PCC connectivity strength is associated with more severe AVH is consistent with prior research demonstrating the link between DMN connectivity and SZ symptoms.^10^ Within-DMN connectivity alterations are not unique to SZ; they have been associated with symptom severity in multiple disorders,^44,45^ such as major depression,^46–48^ addiction,^49^ and chronic pain.^50,51^ This may reflect the DMN’s involvement in transdiagnostic processes such as self-referential thinking,^52–54^ rumination,^55^ and domain-general functions such as allostasis^56,57^ and memory.^58^

This study provides evidence of reduced MPFC-STG/MTG connectivity following STG-based NFB. Importantly, we showed that every participant who received real STG-based neurofeedback reduced STG-MPFC connectivity post-NFB. This is promising for treatment since prior research showed that patients experiencing AHs have disrupted connectivity between midline DMN regions and the STG during rest^16,17^ and during auditory processing.^18^ A previous real-time neurofeedback study using language network (i.e., left STG and IFG) activation as the neurofeedback signal showed that upregulation of the language network led to strengthened STG-MPFC connectivity and that downregulation of the language network led to strengthened STG-PCC connectivity in patients with AHs.^22^ Our previous NFB study using DMN-central executive network activation difference as the neurofeedback signal did not find MPFC-STG connectivity change post-neurofeedback at the group level, although individuals with greater reductions in MPFC-STG connectivity exhibited greater reductions in AHs.^21^ To our knowledge, the current study is the first to report decreased functional connectivity between midline DMN regions and auditory cortices after real-time neurofeedback training. This finding is particularly notable for providing the first direct evidence of the the “resting state hypothesis”,^19,59^ which proposes that AHs result when abnormally high resting state interaction between the MPFC and STG causes misattribution of internally generated auditory stimulus as external. In our study, a single session of STG-based neurofeedback was able to dampen the resting state communication between the MPFC and STG, potentially because STG-based neurofeedback more directly affected the AHs circuitry and mindfulness was a strategy effective at engaging the midline regions.

Notably, we also found reduced connectivity between the MPFC and the left IFG, an important node of the language network that activates during AHs^60–62^ and that shares elevated functional connectivity with the STG in AHs.^63^ Interestingly, a previous STG-based neurofeedback study found strengthened STG-IFG connectivity that was associated with a reduction in AHs, which suggested that an upregulation of speech production circuitry may lead to attenuated response to inner speech.^42^ Our finding of reduced MPFC-IFG connectivity offers further evidence that STG-based neurofeedback combined with mindfulness may modulate the circuitry underlying self-monitoring of speech in AVH.

Several questions remain. First, compared to the sham condition, subjects receiving real NFB did not show greater improvement in their AHs or in MPFC-PCC connectivity. It is possible that despite having received sham neurofeedback, participants in the sham condition still gained benefit via mindfulness practice, which gave rise to modulated functional wiring within the DMN as well as attenuated AHs. A larger sample may have better power to detect a group by time interaction in AHs and MPFC-PCC connectivity. Further, we did not find any association between change in STG-MPFC connectivity and change in AVH, although this lack of finding may be due to the small sample size (*n* = 12 in the real condition) or differential post-NFB assessment timepoints for resting state (immediately post-NFB) and AHs (several weeks post-NFB). A larger sample combined with a longitudinal approach sampling both resting state and AHs at several time points post-NFB may help elucidate the trajectories of neural change and symptom improvement. Future directions for NFB research also include personalization of the neurofeedback training in terms of session length and number of sessions.

In conclusion, in this sham-controlled randomized trial of real-time fMRI neurofeedback, we show additional evidence for a positive correlation between DMN connectivity and AHs severity at baseline, and also demonstrate that STG-based NFB can reduce DMN-STG connectivity in schizophrenia patients who have moderate to severe AHs.

## Acknowledgements

This work was supported by the National Institutes of Health (Grant No. R61MH113751 to MAN and SWG). We thank all the participants who took part in our study and the staff at Massachusetts Institute of Technology, Northeastern University Biomedical Imaging Center, McLean Hospital, and Veterans Affairs for assistance with data collection.

## Notes

### Competing Interest Statement

The authors have declared no competing interest.

