## Supplement for "Real-time fMRI neurofeedback reduces default mode network and auditory cortex functional connectivity in schizophrenia"

**Supplementary Figures** **3**
Figure S1. CONSORT diagram. 3
Figure S2. Auditory hallucination changes post-NFB. 4
Figure S3. Pre and post MPFC connectivity maps. 5
Figure S4. STG connectivity changes post-NFB. 6
**Supplementary Methods** **7**
MRI Acquisition 7
**References** **7**

### Supplementary Figures

**Figure S1. CONSORT diagram.**

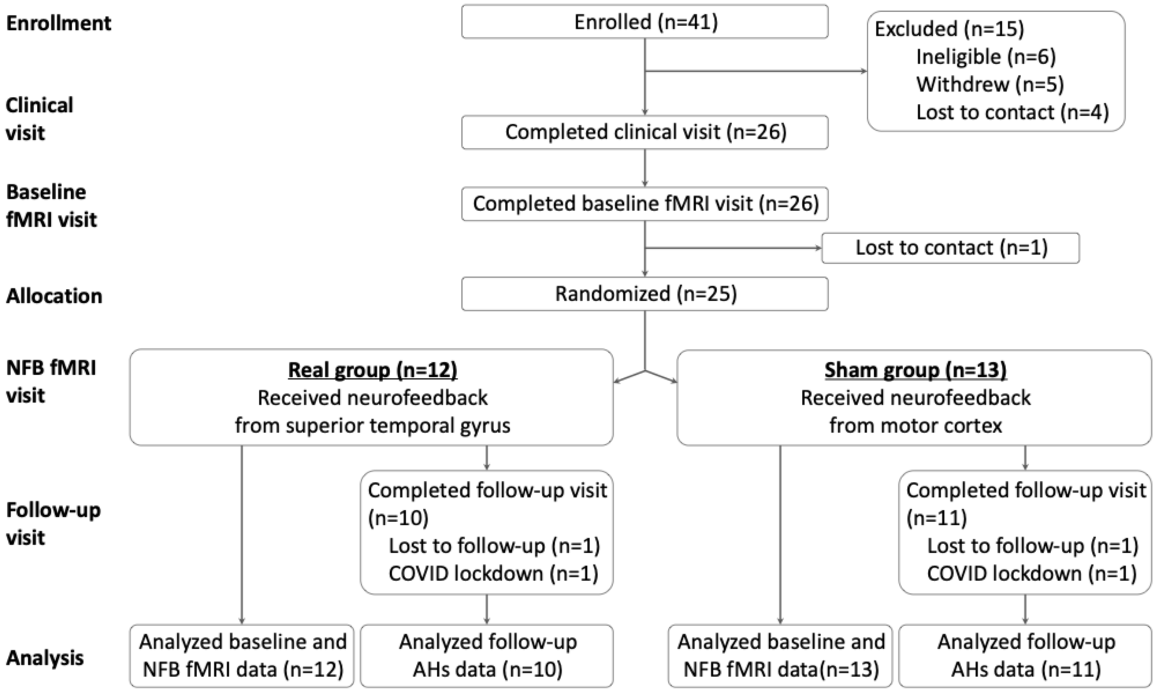


Figure S1. Consolidated Standard of Reporting Trials (CONSORT) diagram.[1] Note that the CONSORT diagram relates to data presented in the current paper, which is a subset of a multi-visit study. Information of the full study can be found on clinicaltrials.gov (NCT03504579) and in other publications.[2,3] AHs: auditory verbal hallucinations; rsfMRI: resting state functional magnetic resonance imaging.

**Figure S2. Auditory hallucination changes post-NFB.**

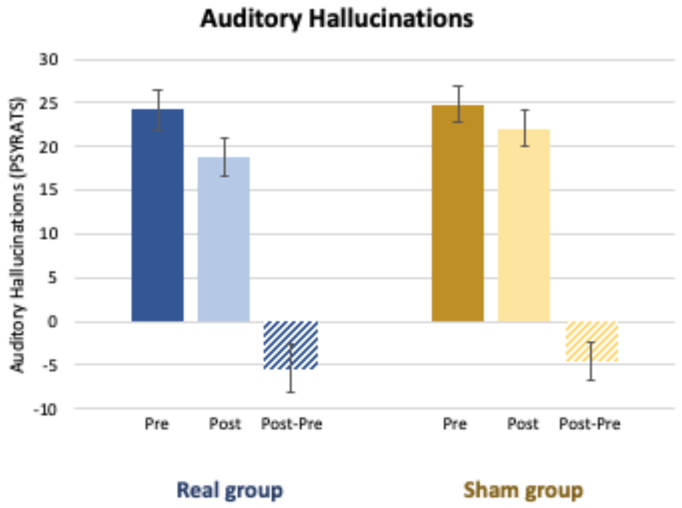


**Figure S2.** Auditory hallucination severity, assessed via the PSYRATS-AH,[4] showed a decrease in both real and sham groups. Error bars indicate one standard error of the mean.

**Figure S3. Pre and post MPFC connectivity maps.**

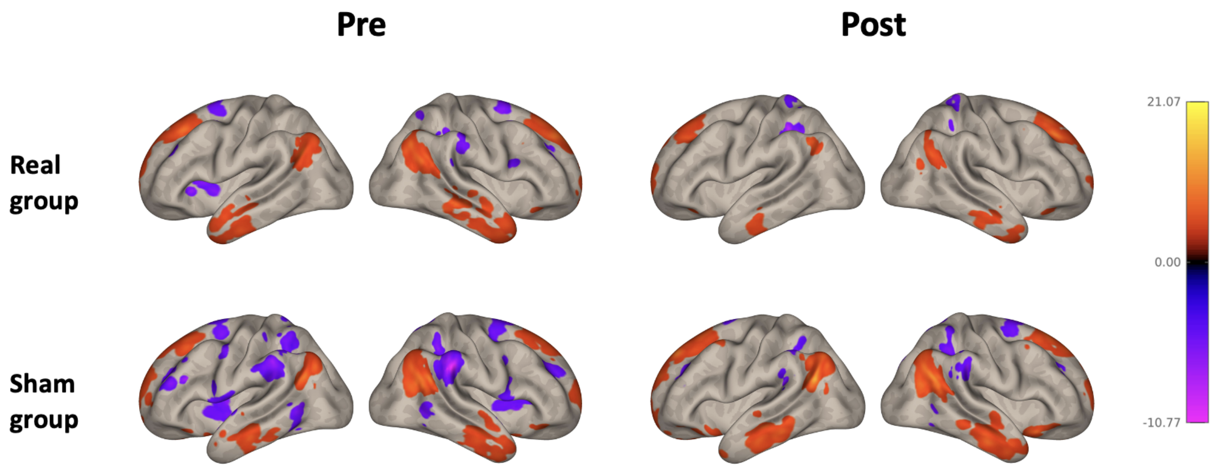


**Figure S3.** Pre and post MPFC connectivity maps are displayed at $p < 0.001$ (uncorrected). Color bar ranges reflect minimum and maximum $t$ values of all maps.

**Figure S4. STG connectivity changes post-NFB.**

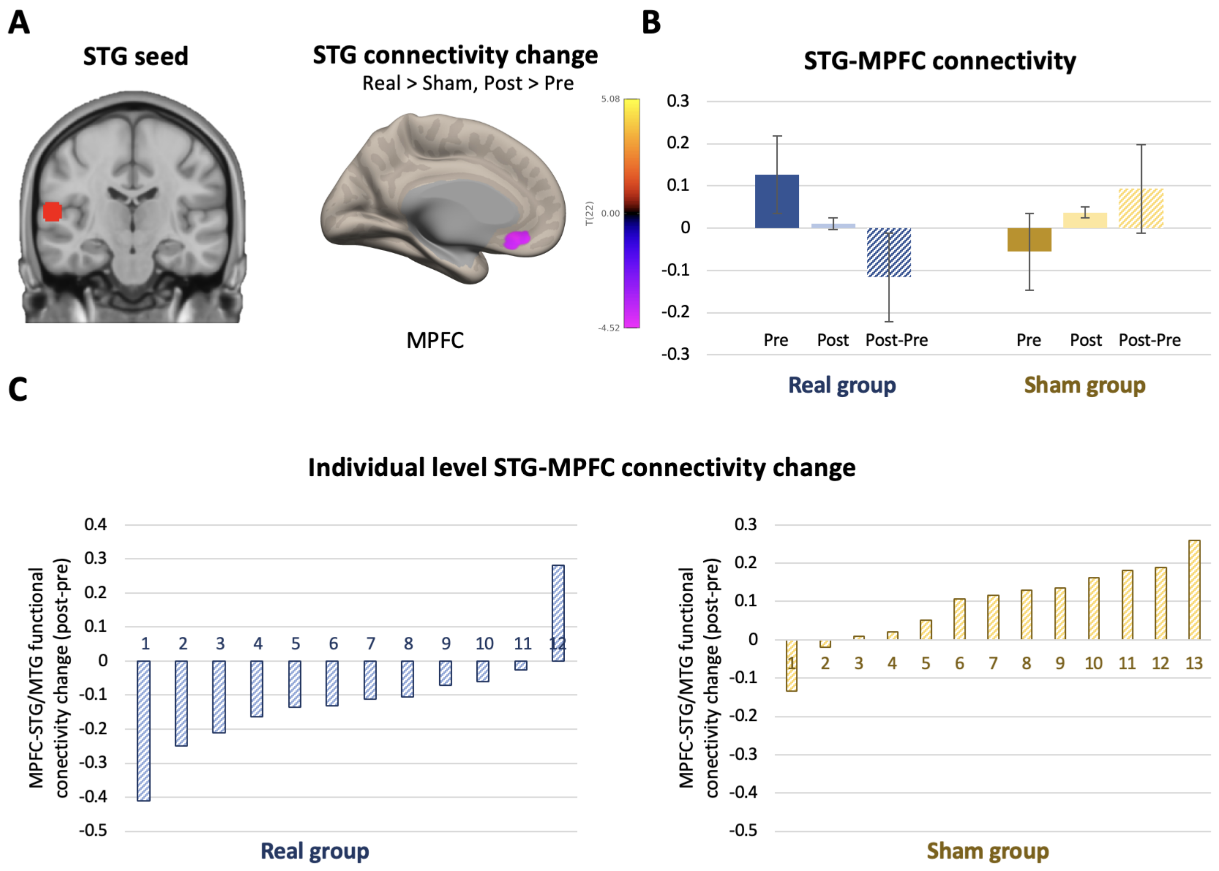


**Figure S4.** One session of NFB reduced STG functional connectivity. **A)** We used an *a priori* STG seed that showed reduced activity post-NFB in a prior study.[5] An ANOVA test revealed that after NFB, there was significantly more reduced connectivity between STG seed and the MPFC ($p < 0.001$, uncorrected) in participants receiving real vs. sham NFB. Color bar ranges reflect minimum and maximum $t$ values in the maps. **B)** The bar plot displays STG-MPFC connectivity pre-NFB and post-NFB, as well as their difference. Error bars indicate one standard error of the mean. **C)** Reduced MPFC-STG/MTG connectivity was found in all but one participant assigned to the real group ($n = 12$) and the opposite pattern of increased MPFC-STG/MTG connectivity was observed in all but one participant assigned to the sham group ($n = 13$).

**Figure S5. STG connectivity changes post-NFB.**

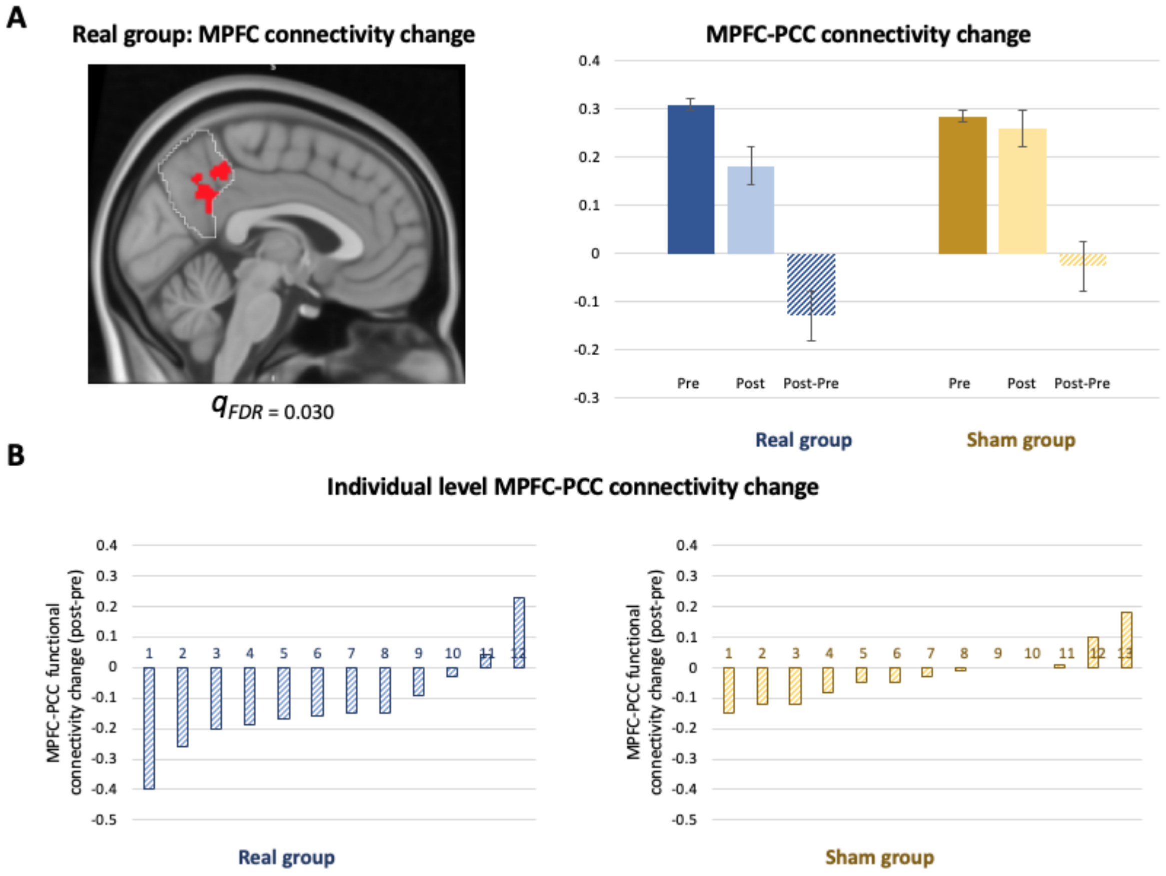


**Figure S5.** One session of NFB reduced MPFC-PCC functional connectivity. **A)** A paired-sample t-test revealed that after NFB, there was significantly more reduced connectivity between MPFC seed and a PCC cluster (small-volume correction using the PCC network node from the CONN toolbox) in participants receiving real. The bar plot displays MPFC-PCC connectivity pre-NFB and post-NFB, as well as their difference. Error bars indicate one standard error of the mean. **B)** Reduced MPFC-PCC connectivity was found in the majority of participants in both groups.

**Supplementary Methods**

**MRI Acquisition**

*MIT Trio* (*n = 7).* Scans were performed using a 32-channel head coil. Structural scans wereacquired using a T1-weighted MPRAGE pulse sequence (1 mm isotropic voxel size, TR = 2530 ms, TE = 1.61 ms, FA = 7°). For functional images, the BOLD signal was measured using a T2* weighted gradient-echo, echo-planar imaging (EPI) pulse sequence (3.5 mm isotropic voxel size, TR = 1090 ms, TE = 30 ms, FA = 90°). Each neurofeedback run lasted 2 minutes and 30 seconds. Immediately before and after NFB, two rs-fMRI scans (5 minutes each) were acquired.

*MIT Prisma (n = 4).* Scans were performed using a 32-channel head coil. Structural scans were acquired using a T1-weighted MPRAGE pulse sequence (1 mm isotropic voxel size, TR = 2530 ms, TE = 1.7 ms, FA = 7°). For functional images, the BOLD signal was measured using a T2* weighted gradient-echo, echo-planar imaging (EPI) pulse sequence (2 mm isotropic voxel size, TR = 1200 ms, TE = 30 ms, FA = 72°). Each neurofeedback run lasted 2 minutes and 30 seconds. Immediately before and after NFB, two rs-fMRI scans (5 minutes each) were acquired.

*Northeastern Prisma (n = 14).* Scans were performed using a 64-channel head coil. Structural scans were acquired using a T1-weighted MPRAGE pulse sequence (0.8 mm isotropic voxel size, TR = 2530 ms, TE = 1.7 ms, FA = 7°). Functional EPI sequence was identical to MIT Prisma.

Ajunwa, C.C., Greene, K.D., Lee, Y., Nestor, P., Kriksciun, M.S., Hammoud, J., Ford, N., Whitfield-Gabrieli, S., Shinn, A.K., Niznikiewicz, M.A. Real-time fMRI neurofeedback modulates auditory cortex activity and connectivity in schizophrenia patients with auditory hallucinations: A controlled study.

4. Drake, R., Haddock, G., Tarrier, N., Bentall, R. & Lewis, S. The Psychotic Symptom Rating Scales (PSYRATS): their usefulness and properties in first episode psychosis. *Schizophr. Res.* **89**, 119–122 (2007).

5. Okano, K. *et al.* Real-time fMRI feedback impacts brain activation, results in auditory hallucinations reduction: Part 1: Superior temporal gyrus -Preliminary evidence. *Psychiatry Res.* **286**, 112862 (2020).
